# Scalable, FACS-Free Genome-Wide Phenotypic Screening

**DOI:** 10.1101/612887

**Authors:** Barbara Mair, Peter M. Aldridge, Randy S. Atwal, Sanna N. Masud, Meng Zhang, David Philpott, Amy H.Y. Tong, Edward H. Sargent, Stéphane Angers, Jason Moffat, Shana O. Kelley

## Abstract

Genome-scale functional genetic screens can identify key regulators of a phenotype of interest, such as determinants of protein expression or modification. Here, we present a rapid, high-throughput approach to phenotypic CRISPR-Cas9 screening. To study factors that modulate the display of CD47 on the cell surface, we processed an entire genome-wide screen containing more than 10^8^ cells in under one hour and maintained high levels of cell viability using a highly scalable cell sorting technology. We robustly identified modulators of CD47 function including QPCTL, an enzyme required for formation of the pyroglutamyl modification at the N-terminus of this protein.

---

Genome-scale genetic screens performed using CRISPR can interrogate determinants of cell viability and are powerful tools for the identification of regulators of phenotypes of interest^1^. Large numbers of cells can be surveyed to isolate rare altered phenotypes using antibodies targeting specific markers of interest. Fluorescence-activated cells sorting (FACS) is the gold standard for sorting and isolation of antibody-labeled cells, but suffers from limited throughput for high-coverage, genome-scale screening applications, resulting in reduced cell viability or necessitating cell fixation due to long sorting times. In addition, it requires advanced, costly instrumentation. Phenotype-based genetic screens^2–8^ can facilitate the identification of regulators of therapeutically relevant proteoforms detected with labelled probes, but are less common compared to proliferation-based screens because of the challenges related to implementation. Thus, rapid and robust selection approaches for targeted capture of live cells are required to realize the full potential of phenotype-based genome-scale screens for functional discovery and further annotation of the human genome.

Here, we describe a high-throughput microfluidic cell sorting platform (HT-miCS) combined with genome-wide CRISPR-Cas9 loss-of-function screening as an unbiased method for identification of modifiers of protein or biomarker expression. In order to identify positive and negative regulators of a biomarker of interest, we designed a microfluidic chip allowing for collection of three subpopulations: a bulk (“medium”) population expressing baseline levels of the target biomarker, and two populations expressing either higher or lower levels (Figure 1A). Cell sorting is facilitated by immunomagnetic labeling using antibodies coupled to magnetic particles. Magnetically-labelled cells are directed to their respective outlets using ferromagnetic guides made of Metglas 2714A, a cobalt-based magnetic alloy, and high-precision manipulation of magnetic drag force and buffer flow. The design of this chip is based on our previous work related to tumour cell sorting^9^; here, we have reengineered the chip to maximize throughput by modifying width and height of the fluidic channel, and the ratio of sample to buffer volume. The HT-miCS chip contains two sets of deflection guides, angled at 5° and 20° relative to the direction of flow to achieve an approximate 10/80/10% distribution of target cells in the low/medium/high outlets (Figure 1B). At an optimized flow rate (Figure S1A), the chip allows processing of 3×10^7^ cells per hour per chip at sorting efficiencies of 73-92% (Figure S1B). In an arrayed setup using 30 parallel chips as an example, this platform can achieve sorting capacities of close to 1 billion cells per hour (Figure 1C).

**Figure 1:**
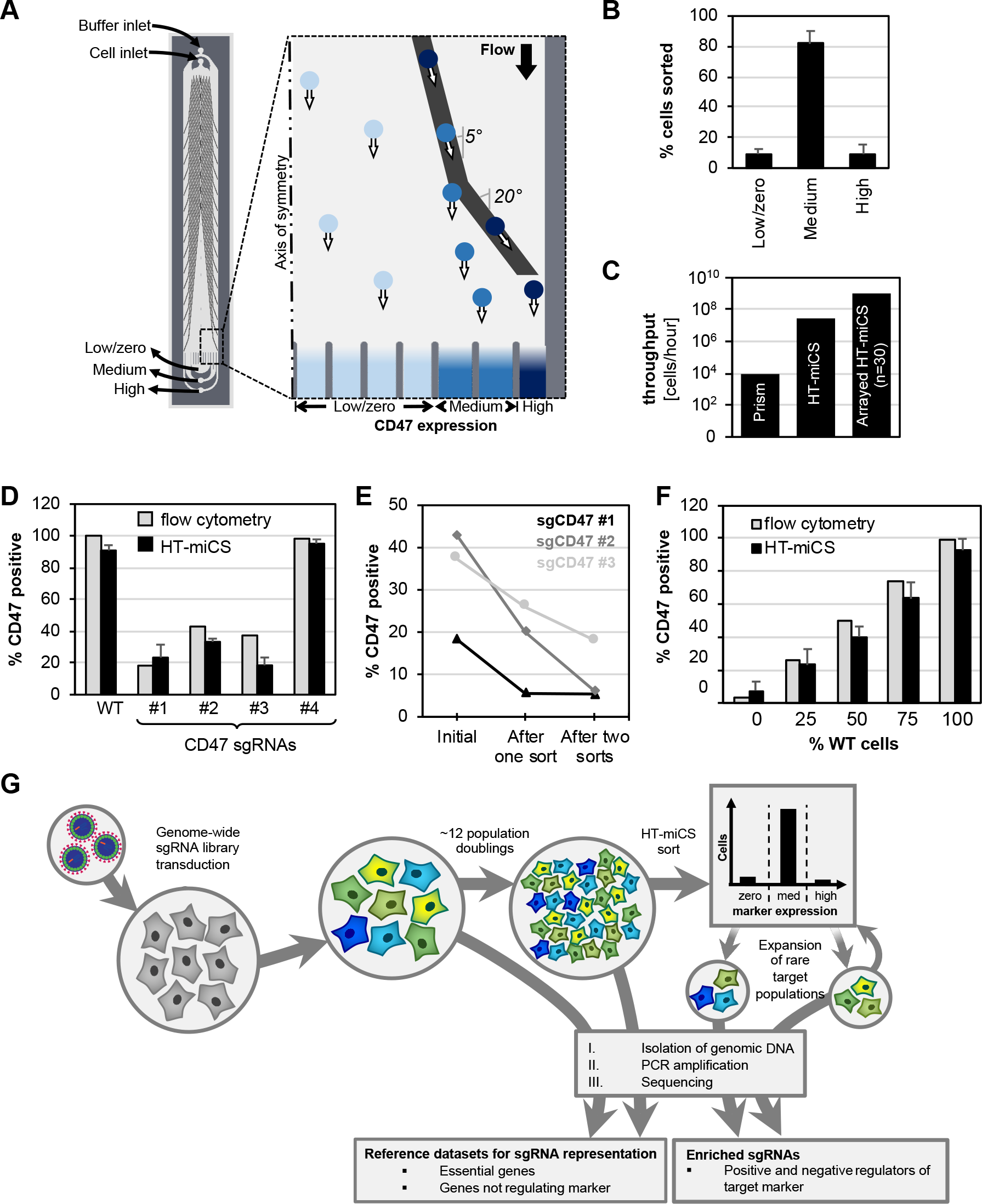
Scalable FACS-free phentotypic screening. **A.** The HT-miCS chip contains two sets of ferromagnetic deflection guides angled at 5° and 20° relative to the direction of flow. A neodymium magnet placed underneath the chip generates a near-uniform magnetic field, and the ferromagnetic guides generate local field amplifications. Magnetically labelled cells, subjected to magnetic and drag forces, will follow the guides as long as the component of drag force acting perpendicular to the guides does not exceed the magnetic trapping force. **B.** Outlet profile of HAP1 cells sorted with the HT-miCS chip, labelled with magnetic beads targeted to CD47 at a sample flow rate of 6ml/hr. Data represent n=3 biological replicates and mean ± SD of triplicates. **C.**Throughput of microfluidic chip sorting. Numbers are calculated based on observed throughput with Prism chip^9^ and HT-miCS (Fig. S1) for 1 and 30 chips. **D.** Comparing HT-miCS performance to flow cytometry. sgRNA #4 does not generate a knock-out of CD47. HT-miCS data represent n=3 technical replicates and mean ± SD of triplicates. Flow cytometry was performed on an aliquot of the same cell pool at 14 days post-transduction. CD47 positive = CD47^med^ + CD47^high^. WT, wild-type. **E.** Sequential sorting of cells transduced with sgRNAs targeting CD47. CD47^low^ cells were expanded after the primary sort, re-sorted 6 days later, expanded and analyzed by flow cytometry. **F.** Sorting defined mixtures of WT and 2-sort-purified pooled CD47 (sgRNA#1) cells. Data represent n=3 technical replicates and mean ± SD of triplicates. Flow cytometry data of same mixtures are shown for comparison. **G.** Schematic of CRISPR screen workflow with two rounds of HT-miCS sorting.

For benchmarking the performance of HT-miCS, we chose an antibody targeting the signal regulatory protein α (SIRPα) binding site on CD47 (CC2C6)^10,11^ for functional profiling of the CD47-SIRPa interaction. CD47 is a widely expressed cell surface protein that acts as a “don’t eat me” signal through inhibitory interactions with SIRPα, a protein expressed on macrophages and other myeloid cells that negatively regulates phagocytosis^12^. CD47 is highly expressed on various tumour types, and blocking the CD47-SIRPα, interaction has been explored as a novel cancer immunotherapy strategy that has shown promising initial results for some cancer types^13–15^.

We transduced HAP1 cells, a near-haploid mammalian cell line widely used for functional genetic screens^16,17^ that robustly expresses CD47^18,19^, with Cas9 and single guide RNAs (sgRNAs) targeting *CD47* and processed the mutants by HT-miCS. In parallel, we performed FACS-based cell sorting. Detection of CD47^low^ cells was comparable between both methods, and recovery (~80%) and cell viability (~90%) after HT-miCS were high, allowing for a secondary sort for further enrichment of CD47^low^ cells (Figures 1D, E). Next, we mixed *CD47* wild-type and knock-out cells in defined ratios and again observed accurate recovery by flow cytometry and HT-miCS (Figure 1F).

For a genome-scale proof-of-concept screen, we then mutagenized HAP1 cells using the Toronto KnockOut version 3.0 (TKOv3) CRISPR library^20–22^. The mutant pool was propagated for ~12 doublings and then subjected to two rounds of microfluidic sorting with recovery, expansion and re-sorting of rare mutants with altered CD47 levels (Figure 1G). SgRNA abundance in each of the sampled populations was determined by deep sequencing and compared between the enriched and unsorted cell populations to identify candidate regulators of CD47 expression. In total, we processed 3×10^8^ target cells on parallelized HT-miCS chips driven by multiple syringe pumps. This arrayed setup allowed for bulk sample preparation, adding no extra processing time with increasing cell numbers, resulting in a net sorting time of ~1 hour (Table S1). For comparison, we performed a FACS-based screen using the same mutagenized cell pool, achieving - at best - sorting rates of 8×10^7^ cells per hour, which is only possible with small, resilient, non-aggregating cell lines (Table S1). We also compared HT-miCS with MACS-based sorting (Table S1), but this approach suffered from poor recovery of viable cells, and detected enrichment of CD47 sgRNAs only in one of three replicates (Table S2). We did not pursue extensive optimization of this approach, leaving open the possibility that it could be made effective, but MACS is likely not amenable to separating cells with subtly different phenotypes.

We calculated normalized z-scores^23^ for the enriched sgRNAs in the CD47^high^ (Tables S3, S4) and CD47^low^ populations (Figure 2A; Tables S3, S4). *CD47* was detected as a strong hit in the CD47^low^ population by both methods (Figures 2A, B; Figure S2; Tables S3, S4). Of note, the three effective *CD47* sgRNAs (Figure 1D) were enriched in the CD47^low^ population in both HT-miCS and FACS, whereas the ineffective sgRNA (#4) was not (Tables S3, S4). In addition to *CD47*, five other top-ranked hits (<20% FDR) overlapped between the FACS and HT-miCS CD47^low^ screens (Figure 2B), of which *QPCTL* was the strongest (Figure S2), in line with a recent report of a gene-trap mutagenesis screen using the same antibody^24^.

**Figure 2:**
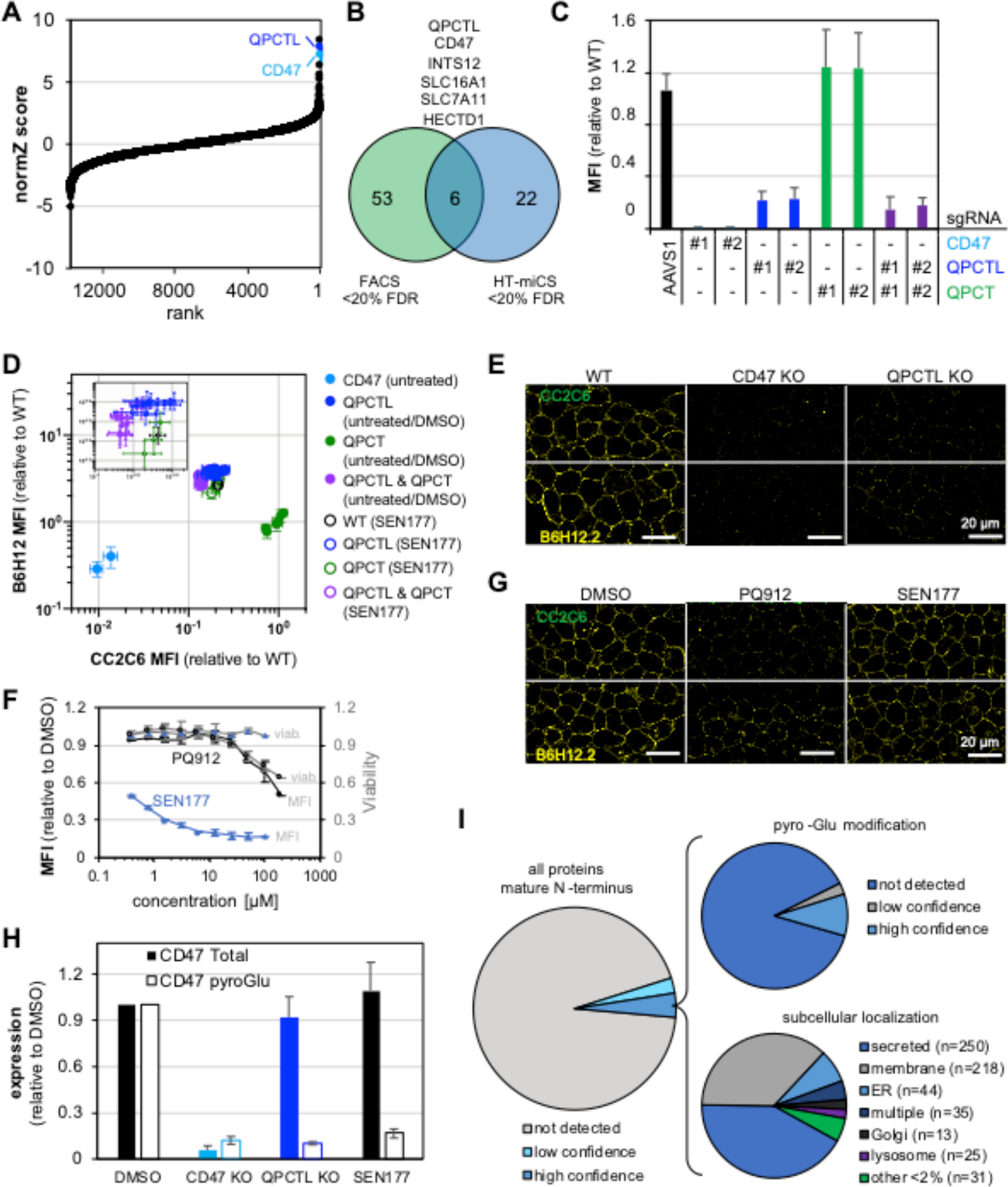
HT-miCS CRISPR screen identifies QPCTL as a modifier of CD47^pyro-Glu^. **A.** Genes targeted by sgRNAs detected in the HT-miCS CD47^low^ screen, ranked by FDR. The normZ score and FDR were calculated using drugZ. **B.** Overlap of hits from the HT-miCS and FACS CD47^low^ screens at <20% FDR. **C.** Cell surface flow cytometry (using the CD47 CC2C6 antibody) of HAP1-Cas9 cells transduced with sgRNAs targeting *CD47*, *QPCTL* and *QPCT* as indicated, 3-6 days post-transduction. AAVS1 sgRNA was used as control. Data represent mean ± SD of the relative median fluorescence intensity (MFI, to wild-type) of n≥2 biological replicates from independent experiments. Purple, double knock-out. See Fig. S3 for full range of sgRNAs and combinations tested. **D.** Cell surface flow cytometry (using the CD47 CC2C6 and B6H12 antibodies) of HAP1 single cell knock-out clones for *CD47*, *QPCTL, QPCT* and *QPCT+QPCTL*. Cells were either left untreated or treated with SEN177 (25µM) or DMSO for 48h. Data representation as in C, n=3 biological replicates from independent experiments, relative to wild-type (untreated) or wild-type + DMSO (treated). Inset shows close-up of *QPCTL* cluster. **E.** Immunofluorescence staining (using the CD47 CC2C6 and B6H12.2 antibodies, upper/green and lower/yellow panels, respectively) and confocal imaging of HAP1 single cell knock-out clones for *CD47* and *QPCTL.* Scale bar = 20µm. **F.** Cell surface flow cytometry of HAP1 cells and viability after SEN177 or PQ912 treatment for 72h. Data represent mean ± SD of MFI or viability relative to DMSO-treated cells from n≥2 biological replicates from independent experiments. **G.** Immunofluorescence images as in E for SEN177- and PQ912-treated (both at 10µM for 72h) HAP1 cells. **H.** PRM-MS assay for endogenous CD47 expression and pyro-Glu modification quantification in HAP1 cells. Data represent mean ± SD of two technical replicates. **I.** *In silico* analysis of potential QPCTL target proteins. The N-terminal amino acid residue of the protein was determined using signal peptide prediction algorithms. Proteins with high-confidence prediction of N-terminal Glu and Gln residues were analyzed for subcellular localization and previous annotation with pyro-Glu modifications. See methods for details.

The *QPCTL* gene encodes glutaminyl-peptide cyclotransferase-like protein (also known as isoQC), a putative Golgi-resident enzyme and paralogue of the secreted glutaminyl-peptide cyclotransferase (QC, encoded by *QPCT*). Both enzymes catalyze the formation of N-terminal pyroglutamate (pyro-Glu) through cyclization of glutamine and glutamate residues^25,26^. Interestingly, an N-terminal pyro-Glu has been detected by crystallographic analysis of the CD47 protein and was suggested to mediate the interaction with SIRPα^27^, but was assumed to arise spontaneously^28^.

We confirmed reduced levels of CD47 pyro-Glu modification (CD47^pyro-Glu^) -as determined by CC2C6 antibody binding -upon transduction of sgRNAs targeting *QPCTL* by cell surface and intracellular flow cytometry, as well as immunofluorescence imaging in HAP1, 293T and KMS11 cell lines (Figures 2C, D, E; Figures S3, S4). sgRNAs targeting *QPCT* did not affect CC2C6 binding, and *QPCT*-*QPCTL* double targeting did not further reduce CD47^pyro-Glu^(Figure 2C). Using a different CD47 antibody (B6H12), we confirmed that overall CD47 protein expression and cell surface localization were not decreased upon transduction with sgRNAs targeting *QPCTL* (Figures 2D, E; Figure S4). We then tested two small molecule inhibitors of QPCTL, SEN177^29^ and PQ912^30–32^, to validate that the enzymatic activity of QPCTL is required for the observed loss of CC2C6 binding, but not B6H12 binding (Figures 2D, F, G; Figures S4, S5). Further, we directly showed reduced pyro-Glu levels on the CD47 N-terminus upon loss of *QPCTL* or SEN177 treatment by targeted mass spectrometry (Figure 2H, Figure S6). Notably, this is the first report of specific detection and direct quantification of endogenous pyro-Glu as a post-translational modification.

An *in silico* prediction of mature human protein sequences (*i.e.* after signal peptide cleavage) revealed ~600 candidates with predicted glutamine or glutamate residues at their N-terminus (Figure 2I). As expected, most candidates are either secreted or membrane proteins, and some have been annotated with pyro-Glu modifications. Apart from CD47, the list of high-confidence candidates contains the angiogenic ribonuclease angiogenin and the exocrine tissue-associated prolactin-inducible protein (PIP), as well as many chemokines and immunoglobulins including the known QPCTL substrates CCL2 and CX3CL1^33,34^. We also identified signaling proteins like Frizzled-7 (*FZD7*) and G-protein-coupled receptor C6A (*GPRC6A*) among potential QPCTL clients, warranting further investigation of the role of QPCTL in post-translational protein modification.

Here, we demonstrate that HT-miCS is a robust platform for functional genetic screening. An arrayed setup of HT-miCS chips can surpass the throughput of traditional FACS-based immunosorting technologies by an order of magnitude, which will expand the applicability of functional, phenotypic screening to fragile cell types. We applied HT-miCS to a genome-wide CRISPR-Cas9 loss-of-function screen, probing genetic regulators of CD47 and yielding overlapping hits with a parallel FACS-based screen. Most notably, we identified and validated QPCTL as a regulator of CD47^pyro-Glu^, corroborating recent findings reporting QPCTL as a potential modifier of CD47-targeted cancer immunotherapy^24^. Supporting this role, high expression of QPCTL has been shown to be a poor prognostic indicator for renal cancer^35^. Interestingly, our screens also identified *SLC16A1* and *SLC7A11*, which have been implicated in (free) pyro-Glu metabolism as transporters of pyro-Glu itself^36^ or glutamine, respectively. Furthermore, QPCTL could have additional substrate proteins that are involved in multiple biological, physiological and pathological processes.

In theory, our HT-miCS platform is also suitable for identifying negative regulators of a biomarker of interest, such as in the CD47^high^ population of our screen (Table S3), or negative regulators of markers with low expression levels. In summary, HT-miCS is a robust, flexible, parallelizable and customizable high-throughput cell sorting platform particularly suitable for genome-scale screening applications.

## ACKNOWLEDGEMENTS

We thank members of the Kelley, Moffat, Angers and Charlie Boone and Brenda Andrews labs for helpful discussions; Katie Chan for TKOv3 library virus preparation; Matej Usaj for help with data analysis; Patricia Mero for administrative assistance; Jelena Tomic for help with tissue culture, and Ethan Cohen for technical assistance. We are grateful to Dionne White and Joanna Warzyszynska for flow cytometry assistance. We thank the the Centre for Applied Genomics (TCAG) at the Hospital for Sick Children (SickKids) for sequencing. This work was supported by grants from the Canadian Institutes for Health Research (J.M., S.O.K.) and Medicine by Design (S.O.K., J.M.). J.M. is a Canada Research Chair in Functional Genomics.

## AUTHOR CONTRIBUTIONS

B.M. and P.M.A. performed most experiments and analyzed data, with help from R.S.A, S.N.M. and D.P., M. Z. and R.S.A. developed the mass spectrometry assay. A.H.Y.T. helped with screen sequencing and data analysis. B.M., P.M.A., R.S.A., J.M. and S.O.K. wrote the manuscript. B.M., P.M.A., E.H.S., S.A., J.M. and S.O.K. designed the study. S.A., J.M. and S.O.K. supervised the study.

---

**Figure S1:**
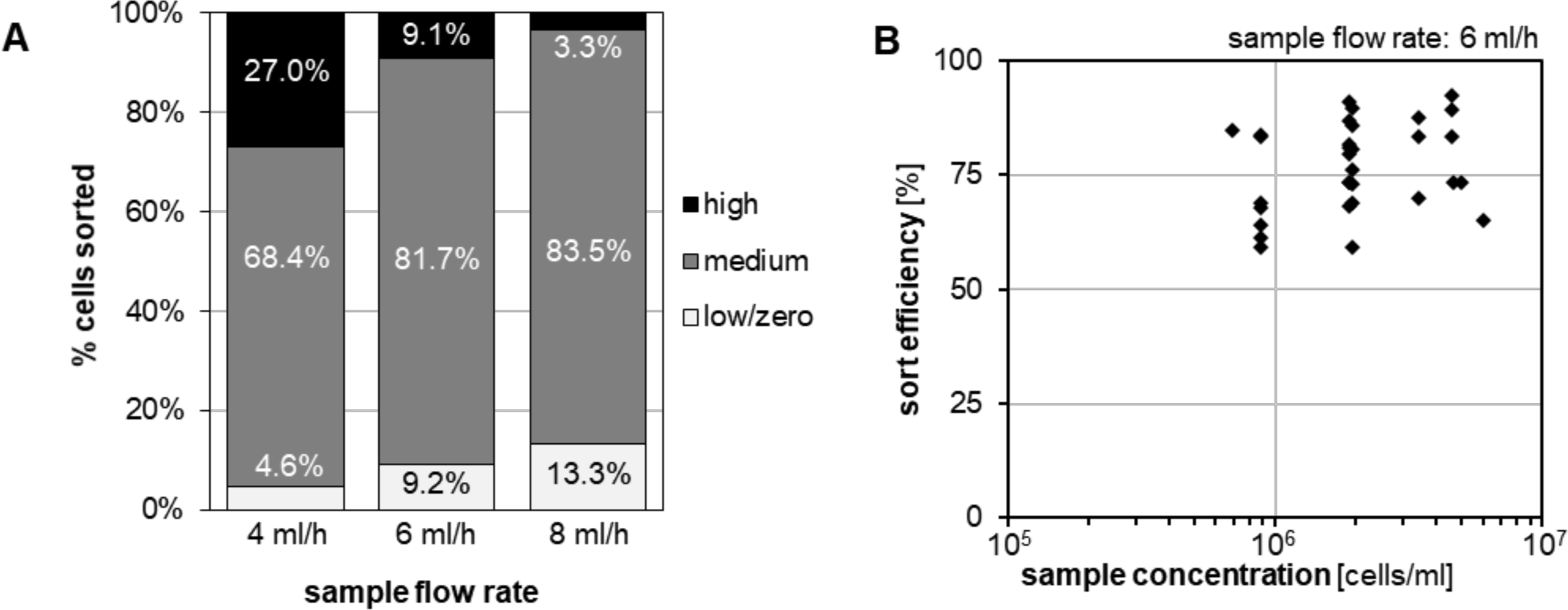
HT-miCS chip optimization. **A.** Flow rate optimization to achieve 10/80/10% distribution into low/zero, medium and high outlets. HAP1 cells were labelled with magnetic beads targeted to CD47. Data represent the mean of n=3 biological replicates. **B.** Throughput optimization to determine maximum sample concentration at optimal flow rate (6 ml/h) and sort efficiency. Sort efficiency was calculated as the percentage of sorted cells directed out of the low/zero outlet. Data represent individual measurements performed using different chips and biological samples.

**Figure S2:**
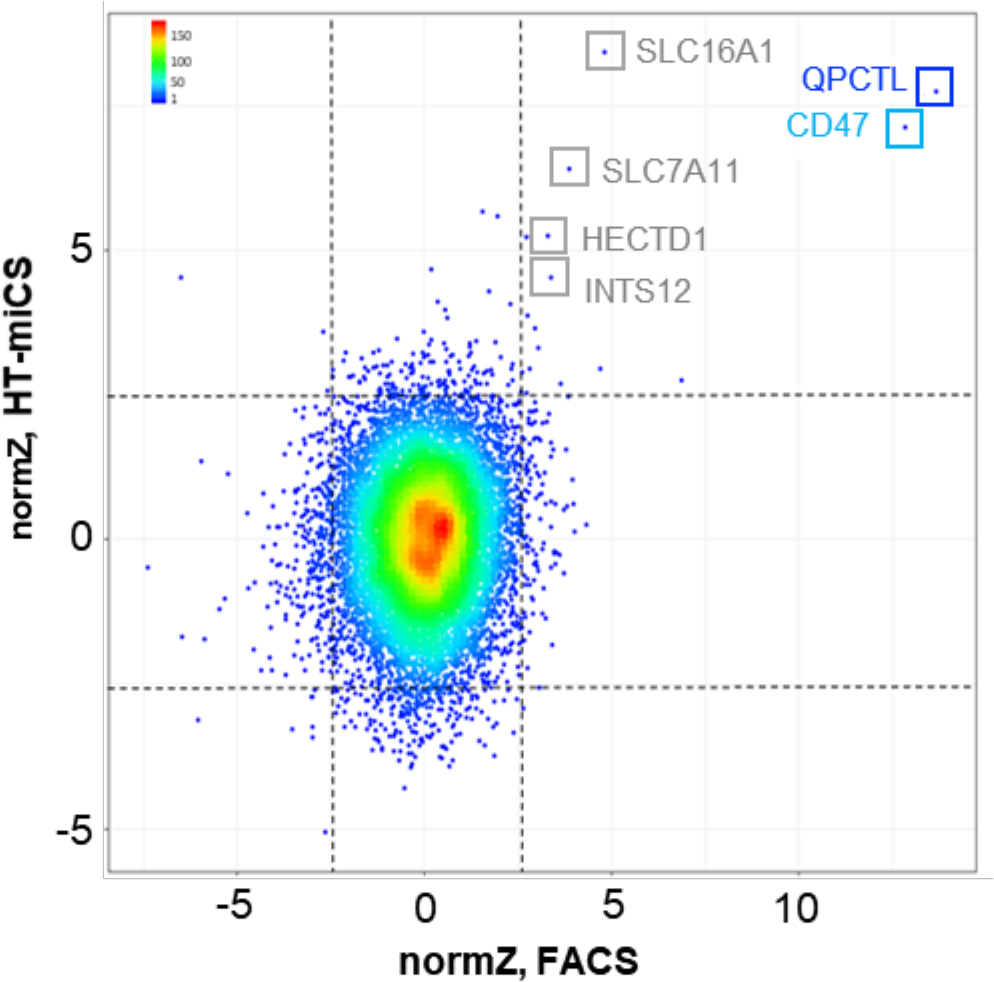
Comparison of HT-miCS and FACS screen results for CD47^low^. Displayed are normZ scores calculated using the drugZ algorithm, coloured by density. Dashed lines represent the top and bottom 90th percentile of the data. Overlapping hits at <20% FDR are highlighted.

**Figure S3:**
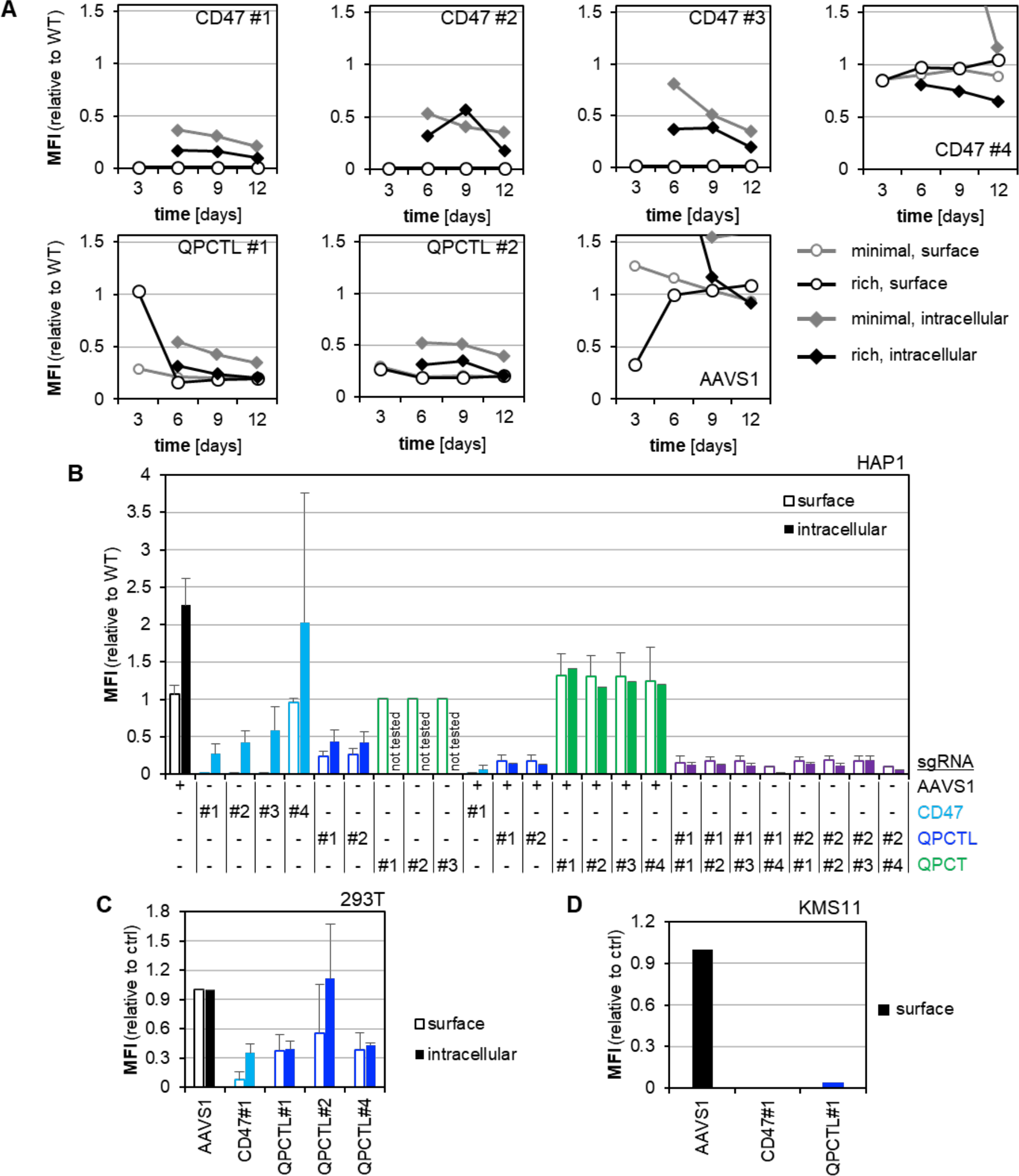
CD47^pyro-Glu^ levels are reduced upon loss of QPCTL. **A.** Cell surface and intracellular flow cytometry (CC2C6 antibody) of HAP1-Cas9 cells transduced with sgRNAs targeting *CD47*, *QPCTL* and *QPCT* as indicated, 3-12 days post-selection in minimal or rich medium. AAVS1 sgRNA was used as control. Data represent the relative median fluorescence intensity (MFI, to wild-type). CD47 #4 and AAVS1 values >1.56 were not plotted. See methods for details. **B.** As in A, 3-6 days post-transduction. Mean ± SD relative MFI of n≥2 biological replicates from independent experiments. **C.** As in B, for 293T cells 5-12 days post-selection. n=2 biological replicates from independent experiments. Data are displayed relative to AAVS1 control. **D.** As in C, for KMS11 cells 25 days post-selection. One representative experiment.

**Figure S4:**
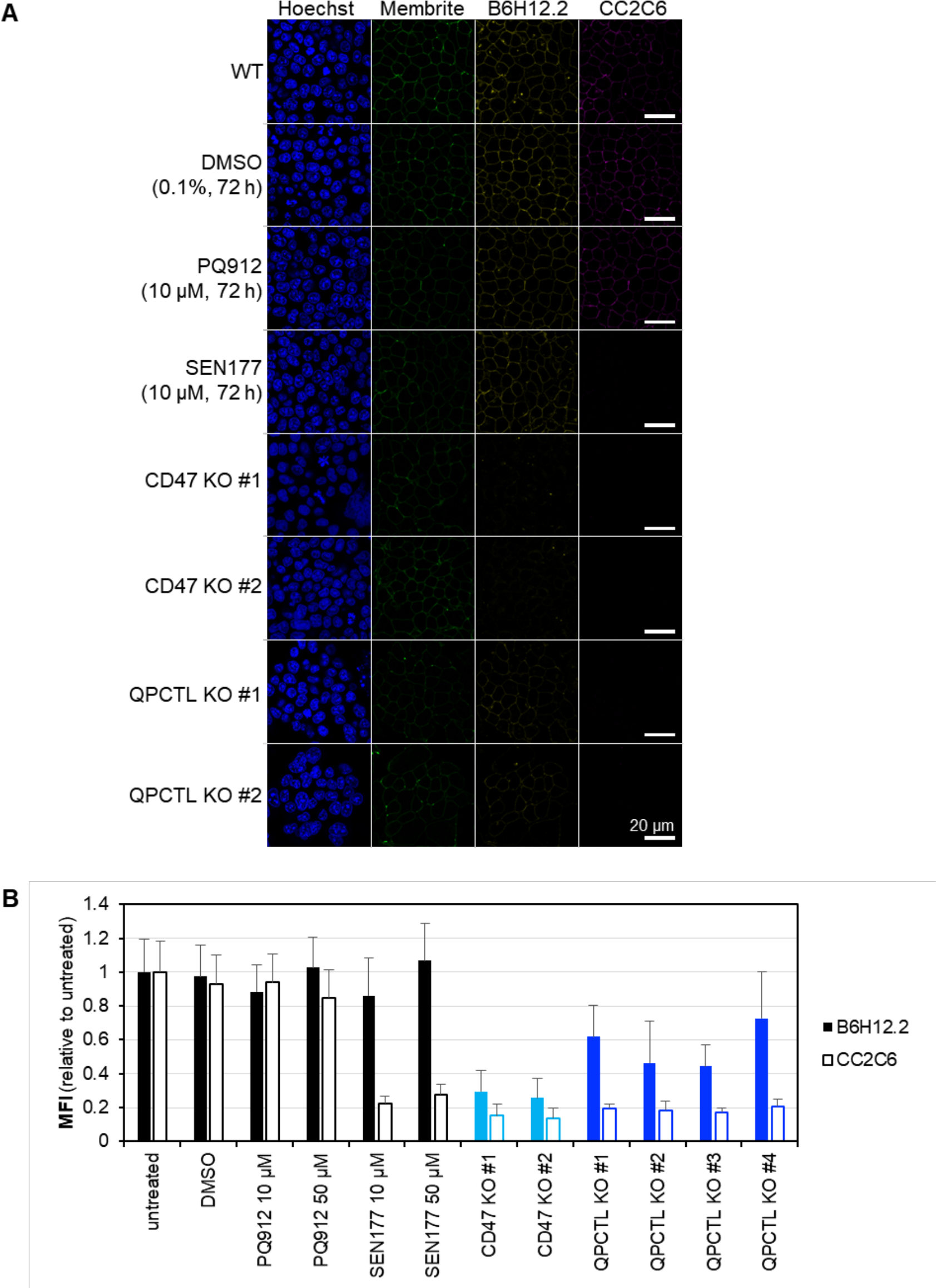
Full range and quantification of immunofluorescence images in Fig. 2E, G. **A.** Immunofluorescence staining (using the CD47 CC2C6 and B6H12.2 antibodies, and Membrite and Hoechst for membrane and nuclear staining as indicated) and confocal imaging of HAP1 single cell knock-out clones for *CD47* and *QPCTL* (*QPCTL* KO #3 and #4 not shown). Scale bar = 20µm. **B.** Quantification of MFI of images (B6H12.2 and CC2C6 only) using a custom ImageJ macro. Data represent the mean ± SD of n=9 technical replicates and n=2 biological replicates. See methods for details.

**Figure S5:**
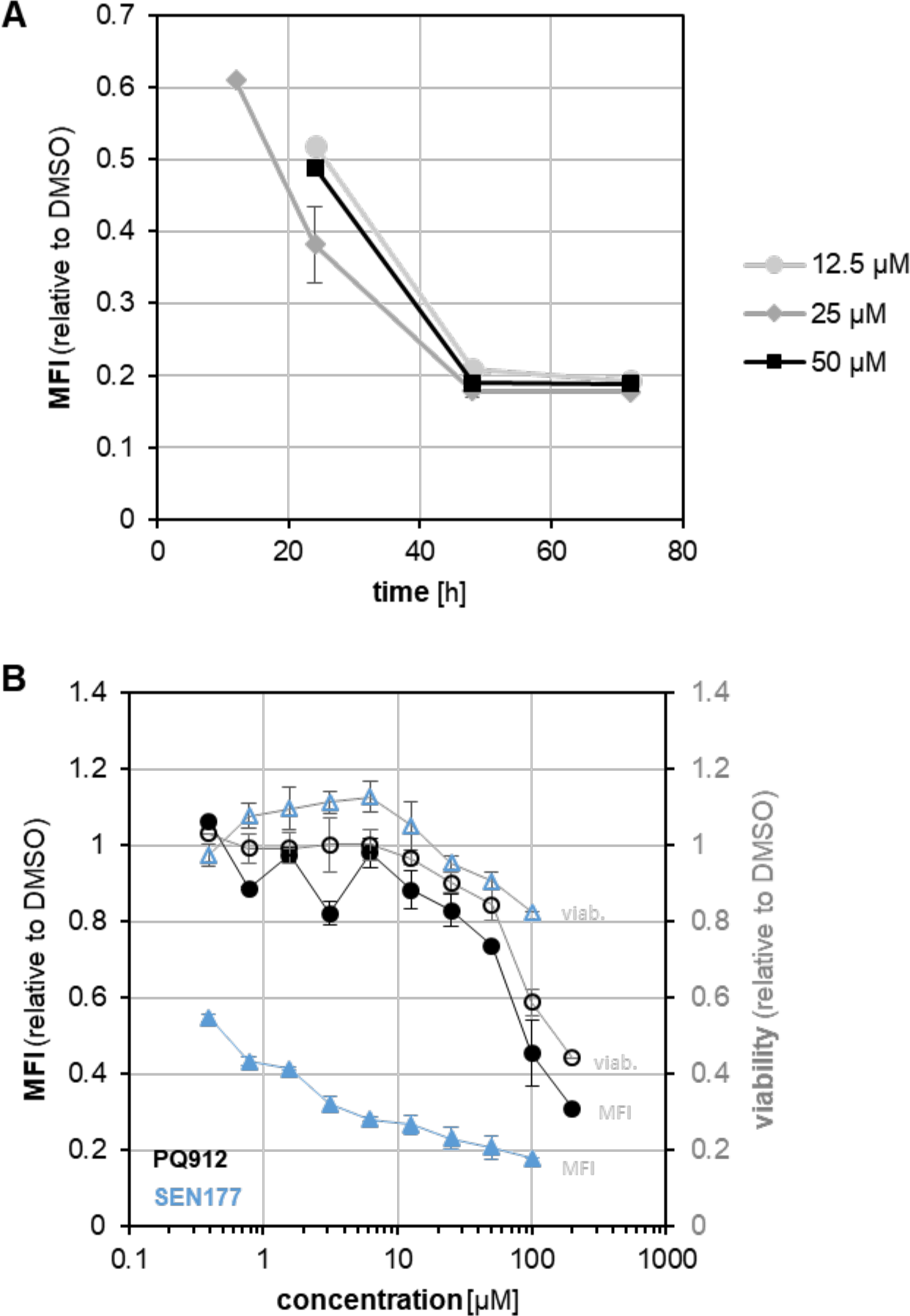
CD47^pyro-Glu^ levels are reduced upon inhibition of QPCTL. **A.** Dose and time dependency of SEN177 treatment of HAP1 cells at indicated concentrations measured by cell surface flow cytometry using the CC2C6 antibody. Data represent the mean ± SD relative median fluorescence intensity (MFI, to DMSO control) of n=2 biological replicates from independent experiments. **B.** Cell surface flow cytometry of 293T cells as in A and viability after SEN177 or PQ912 treatment for 72h.

**Figure S6:**
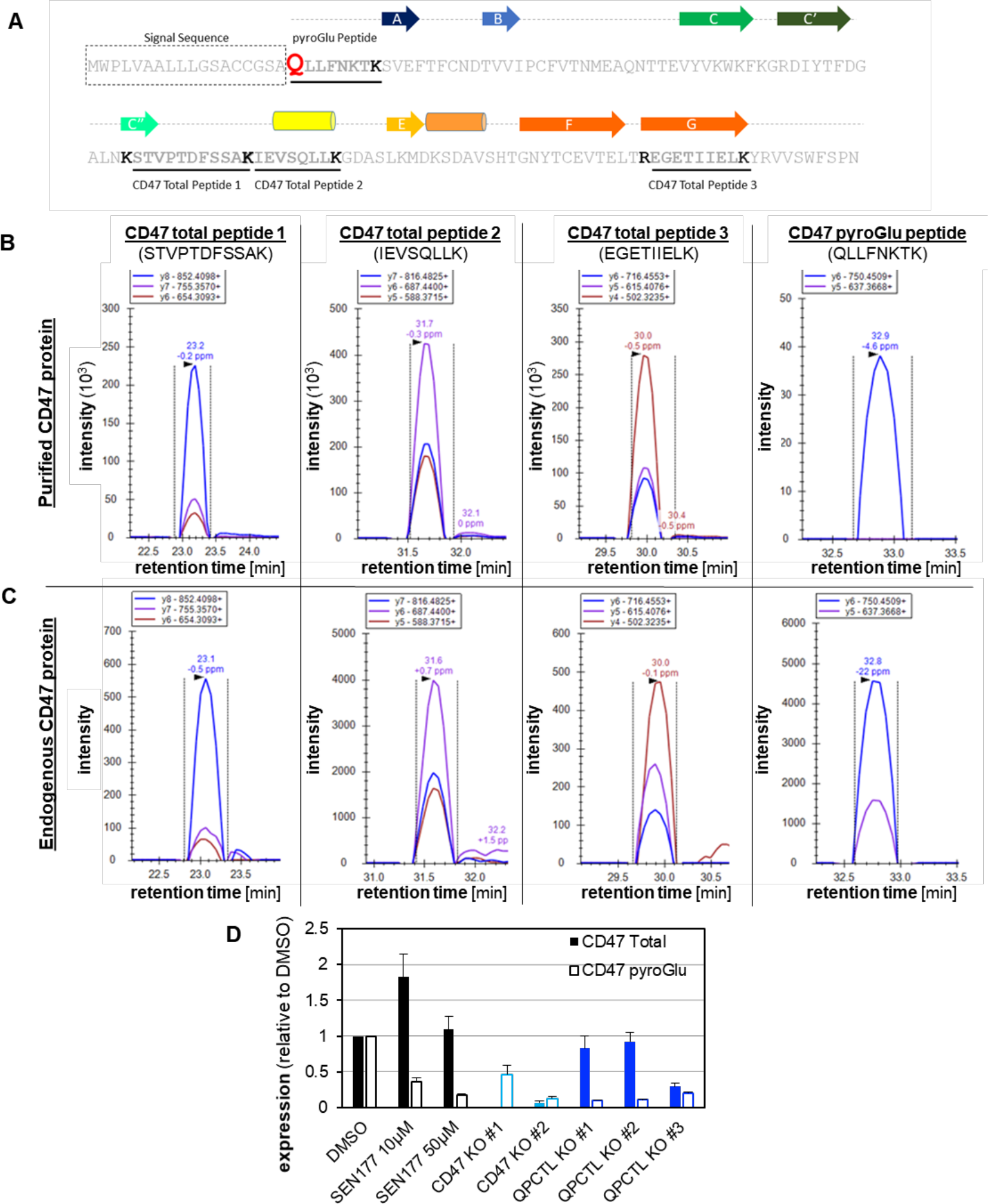
PRM for quantitative MS-based CD47^pyro-Glu^ modification analysis. A. Schematic overlay of the secondary structure (cylinders for alpha helices and arrows for β-sheets) and the primary sequence of the IgSF domain of human CD47 (adapted from Hatherley et. al, Molecular Cell, 2008). Four CD47 tryptic peptides utilized for PRM assays are underlined. Bolded lysine/arginine residues mark the tryptic peptide boundaries. Signal peptide sequence is boxed. The N-terminal glutamine residue (substrate for pyroglutaminyl cyclization) is highlighted in red. **B.** Chromatograms of PRM transitions for the four peptides from extracellular domain of purified recombinant human CD47. **C.** Representative chromatograms of PRM transitions for CD47 peptides from DMSO-treated HAP1 cells for endogenous total CD47 and pyro-Glu level comparisons. **D.** MS-PRM assay for quantitative endogenous total CD47 and pyro-Glu modification level measurements in HAP1 cells. Data represent mean ± SD of two technical replicates.

**Table S1:** Comparison of sorting methods. See methods for details.

|  | HT-miCS | FACS<sup>a</sup> | MACS |
| --- | --- | --- | --- |
| **Outlets** | 3 (low, medium, high) | 2 (custom) | 2 (positive, negative) |
| **Number of markers** | 1 | multiple | 1 |
| **Input CD47 screens** |  |  |  |
| *primary sort* | 118x10<sup>6</sup> cells | 286x10<sup>6</sup> cells | 319x10<sup>6</sup> cells |
| *secondary sort* | 127x10<sup>6</sup> cells | 618x10<sup>6</sup> cells | 29x10<sup>6</sup> cells<sup>b</sup> |
| **Recovery (mean ± SD)** | 84.7±54.93%<sup>c,d</sup> | 50.37±12.79%<sup>e</sup> | 91.9±11.85%<sup>c</sup> |
| **Max. observed sort capacity** | 299x10<sup>6</sup> cells/h<sup>d</sup> | 79x10<sup>6</sup> cells/h<sup>f</sup> | ~106x10<sup>6</sup> cells/h |

<sup>a</sup> BD FACS Aria IIIu
<sup>b</sup> only 2/3 replicates sorted, poor viability/proliferation after primary sort
<sup>c</sup> % of input in all outlets combined, averaged over primary/secondary sorts and all replicates
<sup>d</sup> includes numbers from an unrelated screen performed in parallel, sort time approximated by calculating inlet volume/actual flow rate
<sup>e</sup> % of expected (top and bottom 15% = 30% of input) in all outlets combined, averaged over all timepoints and replicates
<sup>f</sup> replicate C secondary sort CD47<sup>low</sup>

**Tables S2-S4:** Provided as separate files.

## METHODS

### HT-Prism Chip fabrication

Strips of Metglas 2714A were obtained from Metglas and epoxy bonded (Loctite M-31CL, McMaster-Carr) onto 100mm soda lime glass wafers (550µm thick, University Wafer) and left to cure for 24h. Excess epoxy was removed with acetone. The metallic surface was then primed with MCC 80/20 (Microchem) prior to spin coating with S1811 positive photoresist (Microchem). The positive resist was photolithographically patterned, and then the exposed Metglas 2714A was etched using a mixture of 3.6% HCl (Sigma), 14.3% H_12_O_2_ (Sigma) and 82.1% H_2_O. After stripping the remaining photoresist and priming the surface with OmniCoat to improve adhesion, the ferromagnetic guides were encapsulated with a layer of SU-8 3010 (Microchem), and then microfluidic channel features were patterned with a 100 µm layer of SU-8 3050 (Microchem). Each chip was then capped with cured PDMS with holes cored for all inlet and outlet ports, following an APTES treatment (Sigma).^37^ To minimize friction and limit cell adhesion to the chip surfaces, every chip was treated with a solution of 1% w/v pluronic F108 (BASF) in DI H_2_O for a minimum of 12h.^38^

### HAP1 cells

HAP1 cells were obtained from Horizon (clone C631, sex: male with lost Y chromosome, RRID: CVCL_Y019) and HAP1-Cas9 cells were generated as described.^20^ For screening, HAP1 cells were cultured in “minimal” DMEM without sodium pyruvate, with sodium bicarbonate, 1.982g/L glucose and 0.161g/L L-glutamine (Wisent Bioproducts) with 10% FBS (Gibco) and 1% penicillin/streptomycin (ThermoFisher). For all other experiments, cells were cultured in standard medium (IMDM (Gibco) with with 10% FBS (Gibco) and 1% penicillin/streptomycin (ThermoFisher)). The different media conditions do not alter CD47 levels or modification (Fig. S3A). Cells were cultured at 37oC and 5% CO_2_ in humidified incubators, were free of mycoplasma and routinely tested using the MycoAlert Detection Kit (Lonza).

### Other cell lines

HEK293T (CRL-3216, sex: female, RRID: CVCL_0063) were obtained from ATCC and maintained in DMEM (Gibco) with high glucose, L-glutamine and sodium pyruvate, supplemented with 10% FBS (Gibco) and 1% penicillin/streptomycin (ThermoFisher). KMS11 (sex: female, RRID: CVCL_2989) were maintained in RPMI1640 with L-glutamine (Gibco), supplemented with 10% FBS (Gibco) and 1% penicillin/streptomycin (ThermoFisher). Cells were cultured at 37°C and 5% CO_2_ in humidified incubators, were free of mycoplasma and routinely tested using the MycoAlert Detection Kit (Lonza).

### CRISPR sgRNA lentivirus production

pLCKO-TKOv3 plasmid library lentivirus was produced as described.^22^ Briefly, HEK293T cells were seeded at a density of 9×10^6^ cells per 15cm plate and incubated overnight, after which cells were transfected with a mixture of psPAX2 (4.8µg; Addgene #12260), pMDG.2 (3.2µg; Addgene #12259), TKOv3 plasmid library (8µg), and X-treme Gene9 (48µl; Roche) in Opti-MEM (Gibco). 24h after transfection, the medium was changed to DMEM with 1% BSA (Sigma) and 1% penicillin/streptomycin (Gibco). Virus-containing medium was harvested 48h after transfection, centrifuged at 1500rpm for 5min, and stored at −80°C. Functional titers were determined by virus titration on HAP1 cells. 24h after infection, the medium was replaced with puromycin-containing medium (1µg/ml), and cells were incubated for 48h. The multiplicity of infection (MOI) was determined 72h after infection by comparing survival of infected cells to infected unselected and noninfected selected control cells. Lentivirus for individual sgRNA constructs was produced on smaller scale: HEK293T cells were seeded at a density of 0.5×10^6^ per 6-well in low antibiotic growth media (DMEM with 10% FBS (Gibco), 0.1% penicillin/streptomycin) and incubated overnight. Cells were transfected with a mixture of psPAX2 (1800ng), pMDG.2 (200ng), sgRNAs in pLCKO or pLCV2^20,21^ and X-treme Gene9 (12µl) in Opti-MEM. 24h following transfection, medium was changed to serum-free, high BSA growth medium as above. Virus-containing medium was harvested 48h after transfection, centrifuged at 1500rpm for 5min, and stored at −80°C. Functional titers in cells for validation experiments were determined by virus titration.

### Generation of *CD47*, *QPCT* and *QPCTL* knock-out cells

For transduction experiments using cell pools, single-stranded sgRNA oligos were annealed using T4 PNK in T4 ligation buffer (both NEB), and ligated into digested (BsmBI; NEB), phosphatase-treated (rSAP or CIP, NEB) and gel-purified modified pLCKO (for HAP1-Cas9) or pLCV2 (for HAP1, 293T and KMS11) backbones^20,21^ using T4 DNA ligase. All plasmids were Sanger sequence verified and virus was prepared and titered as decribed above. HAP1, HAP1-Cas9 cells, 293T cells or KMS11 cells were infected with target and control gRNAs (MOI <1) in presence of 8µg/ml polybrene. 24h post infection, the medium was replaced with fresh medium containing puromycin (1ug/ml for HAP1 and HAP1-Cas9, 1-2ug/ml for 293T, 0.5ug/ml for KMS11) and cells were incubated for 48h. Cells were further cultured in selection-free medium as indicated for individual experiments (typically T3-T6), and passaged every 3-4 days. CD47 levels or modification did not in-or decrease with prolonged passaging up to T12 in HAP1-Cas9 cells (Fig. S3A). For infections with two sgRNAs (HAP1-Cas9 only), the second sgRNA was cloned into pLCKO hygro (Moffat Lab). Selection was carried out in puromycin (1ug/ml)-and hygromycin (800ug/ml)-containing medium for the first 48h, then in hygromycin-only medium for further 4-5 days. For single-cell knock-out clones (HAP1 only), sgRNAs were cloned into modified PX459v2.0 (Moffat Lab, from Addgene #62988, 1kb stuffer sequence added). HAP1 cells were transfected using Lipofectamine 3000 (3.75µl and 2.3µg plasmid per 6-well), selected with puromycin for 48h as above and seeded in limiting dilutions (1 cell/100µl/96-well) 3 days later. Single-cell colonies were expanded, cryo-banked (in standard medium with 10% DMSO and a total of 20% FBS) and analyzed for mutations as described below. An sgRNA targeting the AAVS1 locus was used as negative control.

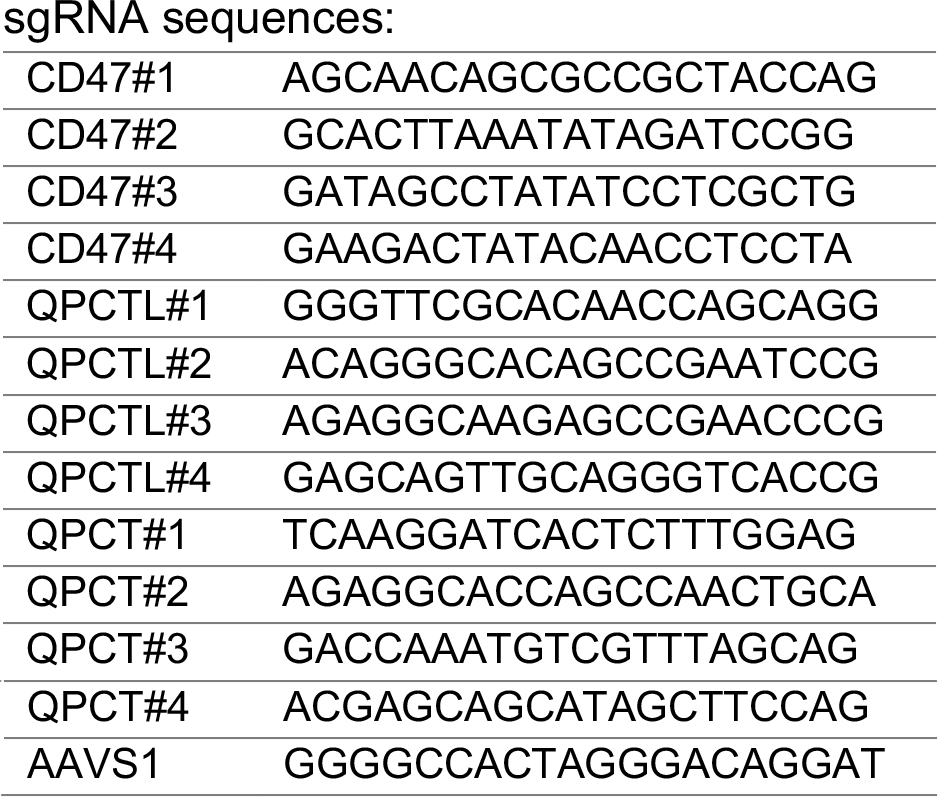

### CRISPR editing analysis

For analysing indels after CRISPR-mediated editing, genomic DNA was isolated using the QIAamp DNA Blood Mini kit (Qiagen) or the Extracta DNA Prep kit (Quantabio). sgRNA target regions (100-200bp up- and 500bp downstream of sgRNA sequence) were amplified by touchdown PCR (−0.5ºC/cycle from 72ºC to 60ºC plus 10 additional cycles at 59ºC) from 50-100ng input DNA, PCR products were purified using the PureLink Quick PCR purification kit (Invitrogen) if necessary and analyzed by Sanger sequencing. Sequences were analyzed to determine the percentage of edited sequences contained in the sample using TIDE^39^ and a “KO-score” using ICE^40^ (Synthego). Non-edited cells amplified with the same primers were used as control samples. A representative analysis of double-sgRNA infected samples from two independent experiments yielded the following editing results for *QPCT* and *QPCTL* target loci (by sgRNA; mean percent edited/KO-score ± SD): QPCTL#1 58.2/NA ± 16.2/N, QPCTL#2 33.23/47 ± 32.96/0, QPCT#1 33.15/56 ± 36.51/0, QPCT#2 80.2/76.4 ± 19.85/2.42, QPCT#3 68.18/36.33 ± 4.29/2.08; AAVS1/CD47#1 (background or off-target, mean ± SD over all *QPCT* and *QPCTL* target loci) 14.4/1.33 ± 14.95/0.58. For single-cell derived clones, TIDE and ICE were used for screening purposes, and mutations were independently confirmed by PCR, Sanger sequencing and sequence alignment as described.

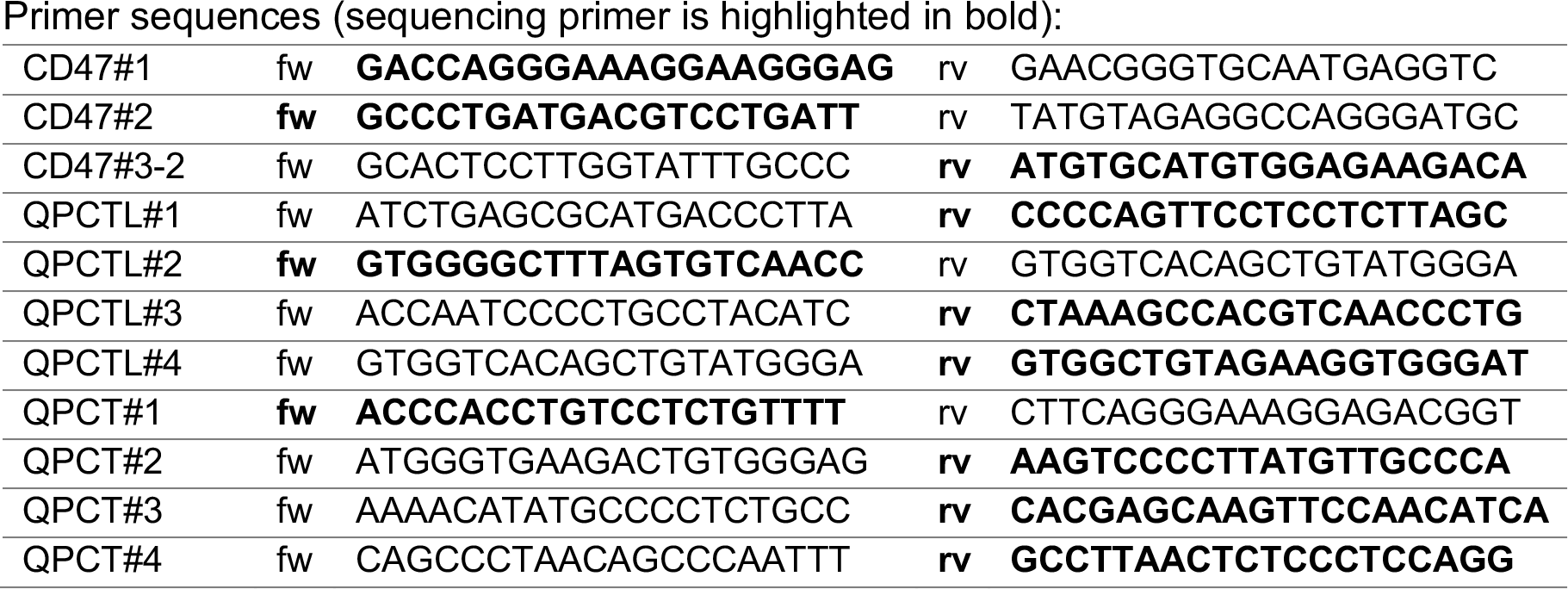

### HT-miCS sorting

Cells were detached with 0.125% trypsin, washed once in PBS with 10% FBS, and resuspended at a concentration of 1×10^7^ cells/ml in a solution of HBSS supplemented with 2% BSA. Cells were labelled for CD47 expression with biotin anti-human CD47 antibody (clone CC2C6, BioLegend, Cat.# 323104, RRID: AB_756134) at 0.0625 µg/100 µl. Excess antibody was removed by washing twice in HBSS with 2% BSA. Cells were resuspended at 1×10^8^ cells/ml, and Anti-Biotin MicroBeads UltraPure were added at a 20% concentration by volume (Cat.# 130-105-637, Miltenyi Biotec) and incubated at room temperature for 30 minutes. Cells were sorted by HT-miCS at a concentration of ~5×10^6^ cells/ml in HBSS with 2% BSA; initial cell concentration was measured using a Countess Automated Cell Counter (Invitrogen). Syringe pumps (Fusion 200, Chemyx), operating in withdrawal mode, were used to drive flow in the HT-miCS chips. Custom 3D-printed mounting hardware allowed for up to five stacks of three syringes (containing 20ml, 10ml and 3ml syringes, Becton Dickinson) to be driven by the same pump. The different cross-sectional areas of the syringes were used to generate different flow rates, corresponding to the width of the low, medium and high outlet channels. Two inlet reservoirs, one containing the sample (cell) solution and one containing a flow focusing buffer stream (HBSS, 2% BSA) were connected to the two inlets of the HT-miCS chip. The pump flow rate was chosen such that the sample flow rate was 6 ml/hr (with a total flow rate of 12 ml/hr). Sorted samples were collected in their respective syringes, and the volume of solution collected in each syringe was measured by weight. A small fraction (100µl) of each sample was collected for cell counting to determine cell concentration after sorting. Each sample was stained with 1µl of Syto24 Green nucleic acid stain (ThermoFisher Scientific) and incubated for 15 minutes at room temperature. Each sample was then loaded onto a 10-chambered microscope slide (Quick Read 3805, Globe Scientific) and cells were counted under fluorescent excitation using a custom counting macro and a Nikon TI Eclipse microscope. Sort efficiency was calculated by dividing the number of cells collected in the medium and high outlets by the total number of cells collected (low, medium and high). Recovery efficiency was defined as the percentage of input cells that were recovered in collected outlet populations. The fractions collected from the low/zero (21%, 24% and 30% of sorted cells for replicates A, B and C, respectively) and high outlets (10%, 18% and 15%) were collected in 15ml falcon tubes on ice. Cells were then spun down, plated in minimal medium and cultured for 6 days prior to secondary sorting (for CD47^low^ fraction from primary sort: 57%, 50% and 35% in low/zero; 1%, 1%, 2% in high). Sorted cells (low/zero and high) were pelleted for genomic DNA extraction. See Table S1 for input cell numbers, recovery and throughput.

### Pooled genome-wide CRISPR screens in HAP1 cells

CRISPR screens in stable HAP1-Cas9 cells were performed essentially as described.^20,21^ Briefly, 150×10^6^ cells were infected with the TKOv3 lentiviral library at an MOI of ~0.3 (>400-fold coverage of the library after selection with puromycin). 24h after infection, medium was changed to puromycin-containing medium (1µg/ml). 72h after infection, 100×10^6^ puromycin-selected cells were cryo-banked, 90×10^6^ cells were split into three replicates of 30×10^6^ cells, passaged every 3-4 days and maintained at 400-fold coverage. 30×10^6^ cells were collected for genomic DNA extraction at T0 after selection and at every passage until day 12 post selection, when sorting was performed. The unsorted T12 sample was used as reference. Genomic DNA extraction, library preparation and sequencing were performed as described below. For the FACS and MACS screens, 90×10^6^ cryo-banked T0 cells were taken in culture, cultured until T12 as above and sorted as described below. The unsorted T12 sample was used as reference. All cell populations tested negative for mycoplasma pre- and post-sorting.

### FACS and MACS sorting

Cells were detached with 0.125% trypsin, counted, and 2×30×10^6^ aliquots were pelleted for genomic DNA extraction. The remaining population was split in half for staining and sorting by FACS and MACS (90-100×10^6^ cells/replicate). For FACS sorting, cells were washed once in PBS and once in Flow buffer (PBS with 2% BSA) and stained with anti-human CD47-APC antibody (clone CC2C6, BioLegend, Cat. #323123/4, RRID: AB_2716202/3) in Flow buffer (20µl antibody/40×10^6^ cells/ml) for 1h rotating at 4ºC in the dark. Cells were washed three times with Sort buffer (PBS wirh 1mM EDTA, 25mM HEPES pH7 and 1% BSA), resuspended in Sort buffer at 40×10^6^ cells/ml, filtered through a 40µm sieve and stained with 7AAD (BioLegend; 50µl/40×10^6^ cells/ml). Small aliquots were taken as single stain controls and stained as above. Sorting was performed on a BD FACS Aria IIIu: 4 laser (405/488/561/633) 15 parameter cuvette based sorter with injection of approximately 40×10^6^ cells/hr. The top and bottom 15% (gated on CD47 histogram of viable 7AAD-negative cells) were collected in 15ml falcon tubes with minimal medium supplemented with 50% FCS on ice. Cells were then spun down, taken up in minimal medium, plated and cultured for 6 days. CD47^low^ and CD47^high^ cells were then detached, counted stained and sorted again as above (approx. 90-120×10^6^ cells/replicate/fraction). Sorting gates were set using unsorted mutagenized cells that had been cultured in parallel. For MACS sorting, a MACS LS column mounted on a MidiMACS (Miltenyi Biotech) was used for sorting. Cells were labelled with magnetic nanobeads targeted to CD47 as described for HT-miCS, suspended in HBSS supplemented with 2% BSA, and sorted through the column. The negative fraction was collected in 15ml falcon tubes with minimal medium supplemented with 50% FCS on ice. Cells were then spun down, taken up in minimal medium, plated and cultured for 7-21 days. Only one replicate could be subjected to a secondary sort due to poor recovery and cell viability. Cells were pelleted after the secondary sort, and gDNA extraction, library preparation, sequencing and data analysis were performed as above. All cell populations tested negative for mycoplasma pre- and post-sorting. See Table S1 for input cell numbers, recovery and throughput.

### Genomic DNA extraction and Illumina sequencing

Genomic DNA was extracted from screen cell pellets using the Wizard Genomic DNA Purification kit (Promega). Sequencing libraries were prepared by amplifying sgRNA inserts via a 2-step PCR using primers that include Illumina TruSeq adaptors with i5 and i7 indices. Resulting libraries were subsequently sequenced on an Illumina HiSeq2500 (RRID: SCR_016383) as described.^21^ Each read was completed with standard primers for dual indexing with Rapid Run V1 reagents. The first 20 cycles of sequencing were dark cycles, or base additions without imaging. The actual 26-bp read begins after the dark cycles and contains two index reads, reading the i7 first, followed by i5 sequences.

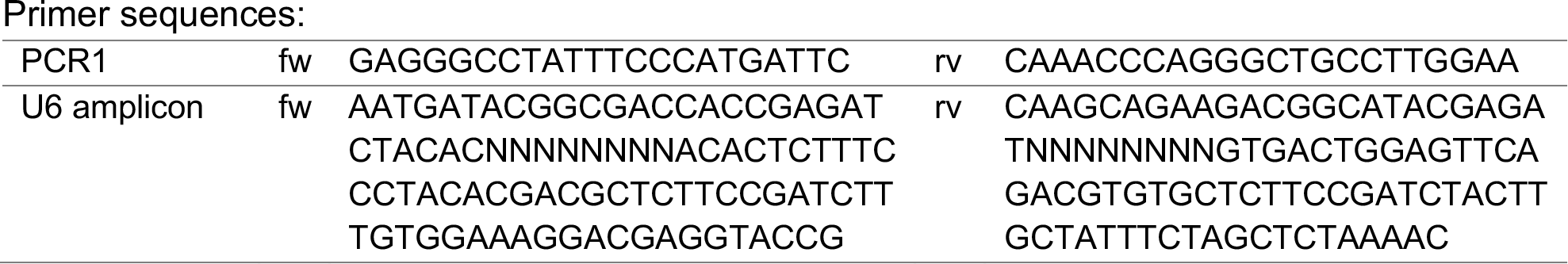

### Screen data processing and quality control

Sample reads were trimmed by locating the first 8bp of the anchors used in the barcoding primers and extracting the flanking 20bp after the anchor was found. We allow a 2bp mismatch for the anchor search. After trimming, a quality control alignment was performed using BOWTIE version 0.12.8 (allowing for max. 2 mismatches, ignoring qualities). For each sample, all available reads were combined from different sequencing runs if applicable, aligned using BOWTIE as described above, and sgRNAs tallied. Read counts for all samples in a screen were combined in a matrix and normalized by dividing each read count by the sum of all read counts in the sample and then multiplying by the expected read number (10 million). Fold change is calculated to a reference sample (T12 unsorted). The calculated fold-changes are then used to generate normZ scores using drugZ.^23^

### Flow cytometry

Cells were dissociated with 0.125% trypsin and washed once in once in Flow buffer (PBS with 2% BSA). For cell-surface analysis, antibody staining was carried out in Flow buffer for 30min on ice at 4°C in the dark. For intracellular antigens, cells were first fixed with 4% PFA (Electron Microscopy Sciences) in PBS for 10min on ice, followed by permeabilization using Flow buffer with 0.1% TritonX-100 (Sigma) for 5min at room temperature. Cells were washed twice with Flow buffer, followed by staining for 30min on ice at 4°C in the dark. Stained cells were washed thrice with Flow buffer and 7-AAD viability dye (3-5µl, BioLegend) was added before quantification. The following antibodies were used for these studies: anti-human CD47-APC (1µl/10^6^ cells in 100µl; clone CC2C6, BioLegend, Cat. #323123/4, RRID: AB_2716202/3), anti-human CD47-FITC (5µl/10^6^ cells in 100µl; clone B6H12, eBioscience, Cat. # 11-0479-4/12, RRID: AB_2043842/3). Stained cells were quantified on an LSRII flow cytometer (BD Biosciences) or an iQue Screener PLUS (IntelliCyt), and data were analyzed using FlowJo software (RRID:SCR_008520). MFI was defined as median fluorescence across the population, and was generally displayed relative to wild-type cells in the same experiment.

### Drug treatments

SEN177 (Sigma) and PQ912 (DC Chemicals) were dissolved in DMSO at 50mM and added to cell medium at indicated concentrations. Cells were incubated for 12-72h as indicated (without refreshing the drugs), and the same volume of DMSO was used as control.

### Immunofluorescence microscopy

Cells were seeded onto poly-D-lysine coated 8-well micro slides (Ibidi). The next day, spent media was removed and cells were washed twice with PBS. Cells were loaded with Membrite^TM^ Fix 488/515 cell surface staining kit (Biotium) per manufacturer’s instructions. Subsequently, cells were washed twice with ice-cold PBS and fixed with 100% Methanol for 10min at −20°C. After two additional washes with PBS, cells were incubated with a blocking solution (5% FBS with 0.05% Tween-20 in PBS) at room temperature for 1h. Primary antibodies (anti-human CD47-APC; clone CC2C6, BioLegend, Cat. #323123/4, RRID: AB_2716202/3; and clone B6H12.2, ThermoFisher Scientific, Cat. #MA5-11895, RRID: AB_11009368) were added at 1:250 dilution in antibody dilution solution (1% FBS with 0.05% Tween-20 in PBS) and stained over night at 4°C. Secondary antibody (Alexa594-conjugated donkey anti-mouse; ThermoFisher Scientific, Cat. #A21203, RRID: AB_2535789) was added at 1:300 dilution in antibody dilution solution and incubated at room temperature for 1h in the dark. Hoechst 33342 (ThermoFisher Scientific) was added at 1:5000 dilution in PBS and stained for 10 min. Finally, cells were washed twice in PBS and imaged on a Leica SP5 700 Confocal Microscope (Zeiss). For mean fluorescence intensity (MFI) measurements, Membrite^TM^ staining intensity was used for automated image segmentation (ImageJ custom macro, available upon request) for operator independent unbiased selection of primary regions of interest (ROI) masks (in the green channel) and the mean fluorescence intensity was measured from the APC and/or Alexa594 channel. In total, 9 random fields of view were imaged per condition. MFI from wild-type untreated HAP1 cells was used to normalize all tested conditions to generate a relative CD47 expression metric.

### Affinity precipitation and LC-MS/MS sample preparation

At 80-90% confluency, cells were harvested from 10cm dishes by scraping and the cell pellets were washed twice in ice-cold PBS. For total cell lysate preparation, cells were lysed in RIPA buffer (ThermoFisher Scientific; supplemented with HALT protease and phosphatase inhibitor (ThermoFisher Scientific)) by incubation for 1h at 4°C with gentle rocking followed by 3 5-second sonication bursts at 10% amplitude. After a 13000xg spin for 30min at 4°C, the supernatant was collected. Total protein concentrations were measured using the BCA Protein Assay kit (ThermoFisher Scientific). For immunoprecipitation of endogenous CD47, either clone B6H12.2 (for total protein; ThermoFisher Scientific, Cat. #MA5-11895, RRID: AB_11009368) or CC2C6 (for pyro-Glu modified protein; BioLegend, Cat. #323123/4, RRID: AB_2716202/3) anti-CD47 antibodies were used with Protein-G Dynabeads^TM^ or Biotin-binder Dynabeads^TM^, respectively, following manufacturer’s instructions (ThermoFisher Scientific). For quantitative N-terminal CD47^pyro-^Glu modification detection, CD47 immunoprecipitates were eluted in 100mM ammonium bicarbonate (pH 7.5) and digested with trypsin (ThermoFisher Scientific) overnight at 37°C to generate tryptic peptides. Peptides were de-salted using PepClean^TM^ C18 spin-columns following manufacturer’s instructions (ThermoFisher Scientific).

### High-performance liquid chromatography (HPLC)

EASY-nLC 1200 (ThermoFisher Scientific) was coupled to the Q-Exactive MS (ThermoFisher Scientific) for peptide separation and detection. An EASY-Spray column (2μm, 100A, 75μmx50cm; ThermoFisher Scientific) was employed for compound separation. Mobile phase A (0.1% formic acid in H_2_0) and mobile phase B (0.1% formic acid in acetonitrile) were used with the following gradient: 0min, 5% B; 50min, 35% B; 55min, 100% B; 60min, 100% B at a flow rate of 225nl/min.

### Q-Exactive – PRM assays

Samples were analyzed using a Q-Exactive HF quadrupole Orbitrap mass spectrometer (ThermoFisher Scientific). 6 peptides were monitored through a PRM acquisition composed of 1 MS1 scan followed by 6 targeted MS/MS scans in high-energy collision dissociation with cycle times of 2.7s. For generating CD47-specific peptides suitable for the PRM assay, we purchased purified extracellular domain of human CD47 (G&P Biosciences) and used trypsin digestion as above to generate tryptic peptide-specific PRM transition profiles. The recombinant purified CD47 extracellular domain is expected to contain very little pyro-Glu modification (arising primarily due to spontaneous conversion). Acquisition was performed in positive ion mode for all 7 peptides. Due to a low number of target peptides, data acquisition was performed in an unscheduled MS/MS assay but retention times were noted.

### Mass-spectrometry data analysis

PRM data acquired by LC-MS/MS was imported into Skyline for peak extraction and peak area calculation for each peptide. The top 3 fragments for each ion were used for quantification. For quantitative comparison across samples, DMSO control samples were used to generate a relative expression metric for both total CD47 protein expression and pyro-Glu modification across multiple conditions tested.

### *In silico* prediction of QPCTL targets

To predict potential candidates for N-terminal pyro-Glu modification, knowledge of the first amino acid of the mature N-terminus (*i.e.* Q or E) is required. Many secretory or membrane proteins, as well as Golgi- and ER-resident proteins, contain signal peptides (SPs), which are proteolytically removed, revealing the mature N-terminus. As SP processing is a major (but not the only) proteolytic maturation event, we decided to use SP prediction tools to derive an approximation of a mature human proteome. FASTA sequences of human proteins were downloaded from Uniprot (filters: evidence at protein level and reviewed) and used as input for SignalP 4.0^41^ (via SecretSanta^42^), signalHSMM^43^ and Phobius^44^. All tools were run from within R. The following parameters were used for SignalP 4.0: version 4.1, organism ‘euk’, run_mode ‘starter’, sensitive TRUE. For signalHSMM, an SP probability cutoff of >0.45 was used. If a signal peptide was detected, the first residue after the predicted cleavage site was used as the new mature N-terminal amino acid, otherwise the original N-terminus was used. Proteins were grouped according to prediction confidence for putative Gln or Glu (Q or E) N-termini (by 0 tools = not detected, 1 = low confidence, 2/3 = high confidence). We then sub-classified the high-confidence group according to subcellular localization (downloaded from Uniprot followed by manual formatting) and previously detected pyro-Glu modification (downloaded from Uniprot, PDB and dbptm, http://dbptm.mbc.nctu.edu.tw/; by 0 sources = not detected, 1 = low confidence, 2/3 = high confidence). Note that annotation with pyro-Glu in PDB does not necessarily predict direct modification due to the presences of multiple entities in a crystal structure. 67/86, 53/71 and 9/24 pyro-Glu annotations are in the high-confidence Q/E N-terminus group for Uniprot, dbptm and PDB, respectively. Localizations occurring at <2% of total were grouped together under “other”.

## REFERENCES

1. Sharma, S. & Petsalaki, E. Application of CRISPR-Cas9 Based Genome-Wide Screening Approaches to Study Cellular Signalling Mechanisms. Int. J. Mol. Sci. 19, (2018).

2. Burr, M. L. et al. CMTM6 maintains the expression of PD-L1 and regulates anti-tumour immunity. Nature 549, 101–105 (2017).

3. Mezzadra, R. et al. Identification of CMTM6 and CMTM4 as PD-L1 protein regulators. Nature 549, 106–110 (2017).

4. Brockmann, M. et al. Genetic wiring maps of single-cell protein states reveal an off-switch for GPCR signalling. Nature 546, 307–311 (2017).

5. Wroblewska, A. et al. Protein Barcodes Enable High-Dimensional Single-Cell CRISPR Screens. Cell 175, 1141–1155.e16 (2018).

6. de Groot, R., Lüthi, J., Lindsay, H., Holtackers, R. & Pelkmans, L. Large-scale image-based profiling of single-cell phenotypes in arrayed CRISPR-Cas9 gene perturbation screens. Mol. Syst. Biol. 14, e8064 (2018).

7. Haney, M. S. et al. Identification of phagocytosis regulators using magnetic genome-wide CRISPR screens. Nat. Genet. 50, 1716–1727 (2018).

8. Parnas, O. et al. A Genome-wide CRISPR Screen in Primary Immune Cells to Dissect Regulatory Networks. Cell 162, 675–686 (2015).

9. Aldridge, P. M. et al. Prismatic Deflection of Live Tumor Cells and Cell Clusters. ACS Nano 12, 12692–12700 (2018).

10. Seiffert, M. et al. Human signal-regulatory protein is expressed on normal, but not on subsets of leukemic myeloid cells and mediates cellular adhesion involving its counterreceptor CD47. Blood 94, 3633–43 (1999).

11. Leclair, P. et al. CD47-ligation induced cell death in T-acute lymphoblastic leukemia. Cell Death Dis. 9, 544 (2018).

12. Matlung, H. L., Szilagyi, K., Barclay, N. A. & van den Berg, T. K. The CD47-SIRPα signaling axis as an innate immune checkpoint in cancer. Immunol. Rev. 276, 145–164 (2017).

13. Weiskopf, K. Cancer immunotherapy targeting the CD47/SIRPα axis. Eur. J. Cancer 76, 100–109 (2017).

14. Advani, R. et al. CD47 Blockade by Hu5F9-G4 and Rituximab in Non-Hodgkin’s Lymphoma. N. Engl. J. Med. 379, 1711–1721 (2018).

15. Kong, F. et al. CD47: a potential immunotherapy target for eliminating cancer cells. Clin. Transl. Oncol. 18, 1051–1055 (2016).

16. Carette, J. E. et al. Ebola virus entry requires the cholesterol transporter Niemann-Pick C1. Nature 477, 340–3 (2011).

17. Bürckstümmer, T. et al. A reversible gene trap collection empowers haploid genetics in human cells. Nat. Methods 10, 965–971 (2013).

18. Lee, S.-E. et al. Proteogenomic Analysis to Identify Missing Proteins from Haploid Cell Lines. Proteomics 18, e1700386 (2018).

19. Paulo, J. A. & Gygi, S. P. Isobaric Tag-Based Protein Profiling of a Nicotine-Treated Alpha7 Nicotinic Receptor-Null Human Haploid Cell Line. Proteomics 18, e1700475 (2018).

20. Hart, T. et al. High-Resolution CRISPR Screens Reveal Fitness Genes and Genotype-Specific Cancer Liabilities. Cell 163, 1515–1526 (2015).

21. Hart, T. et al. Evaluation and Design of Genome-Wide CRISPR/SpCas9 Knockout Screens. G3 7, 2719–2727 (2017).

22. Mair, B. et al. Essential Gene Profiles for Human Pluripotent Stem Cells Identify Uncharacterized Genes and Substrate Dependencies. Cell Rep. 27, 599–615.e12 (2019).

23. Colic, M. et al. Identifying chemogenetic interactions from CRISPR knockout screens with drugZ. bioRxiv 232736 (2019). doi:10.1101/232736

24. Logtenberg, M. E. W. et al. Glutaminyl cyclase is an enzymatic modifier of the CD47-SIRPα axis and a target for cancer immunotherapy. Nat. Med. 25, 612–619 (2019).

25. Cynis, H. et al. Isolation of an Isoenzyme of Human Glutaminyl Cyclase: Retention in the Golgi Complex Suggests Involvement in the Protein Maturation Machinery. J. Mol. Biol. 379 966–980 (2008).

26. Stephan, A. et al. Mammalian glutaminyl cyclases and their isoenzymes have identical enzymatic characteristics. FEBS J. 276, 6522–36 (2009).

27. Hatherley, D. et al. Paired Receptor Specificity Explained by Structures of Signal Regulatory Proteins Alone and Complexed with CD47. Mol. Cell 31, 266–277 (2008).

28. Ho, C. C. M. et al. “Velcro” Engineering of High Affinity CD47 Ectodomain as Signal Regulatory Protein α (SIRPα) Antagonists That Enhance Antibody-dependent Cellular Phagocytosis. J. Biol. Chem. 290, 12650–12663 (2015).

29. Pozzi, C., Di Pisa, F., Benvenuti, M. & Mangani, S. The structure of the human glutaminyl cyclase-SEN177 complex indicates routes for developing new potent inhibitors as possible agents for the treatment of neurological disorders. J. Biol. Inorg. Chem. 23, 1219–1226 (2018).

30. Ramsbeck, D. et al. Structure-activity relationships of benzimidazole-based glutaminyl cyclase inhibitors featuring a heteroaryl scaffold. J. Med. Chem. 56, 6613–25 (2013).

31. Lues, I. et al. A phase 1 study to evaluate the safety and pharmacokinetics of PQ912, a glutaminyl cyclase inhibitor, in healthy subjects. Alzheimer’s Dement. Transl. Res. Clin. Interv. 1, 182–195 (2015).

32. Hoffmann, T. et al. Glutaminyl Cyclase Inhibitor PQ912 Improves Cognition in Mouse Models of Alzheimer’s Disease—Studies on Relation to Effective Target Occupancy. J. Pharmacol. Exp. Ther. 362, 119–130 (2017).

33. Cynis, H. et al. The isoenzyme of glutaminyl cyclase is an important regulator of monocyte infiltration under inflammatory conditions. EMBO Mol. Med. 3, 545–558 (2011).

34. Kehlen, A. et al. N-terminal pyroglutamate formation in CX3CL1 is essential for its full biologic activity. Biosci. Rep. 37, (2017).

35. Uhlen, M. et al. A pathology atlas of the human cancer transcriptome. Science 357 eaan2507 (2017).

36. Sasaki, S., Futagi, Y., Kobayashi, M., Ogura, J. & Iseki, K. Functional Characterization of 5-Oxoproline Transport via SLC16A1/MCT1. J. Biol. Chem. 290, 2303–2311 (2015).

37. Ren, Y. et al. A simple and reliable PDMS and SU-8 irreversible bonding method and its application on a microfluidic-MEA device for neuroscience research. Micromachines 6, 1923–1934 (2015).

38. Luk, V. N., Mo, G. C. & Wheeler, A. R. Pluronic Additives: A Solution to Sticky Problems in Digital Microfluidics. Langmuir 24, 6382–6389 (2008).

39. Brinkman, E. K., Chen, T., Amendola, M. & van Steensel, B. Easy quantitative assessment of genome editing by sequence trace decomposition. Nucleic Acids Res. 42, e168–e168 (2014).

40. Hsiau, T. et al. Inference of CRISPR Edits from Sanger Trace Data. bioRxiv 251082 (2018). doi:10.1101/251082

41. Petersen, T. N., Brunak, S., von Heijne, G. & Nielsen, H. SignalP 4.0: discriminating signal peptides from transmembrane regions. Nat. Methods 8, 785–6 (2011).

42. Gogleva, A., Drost, H.-G. & Schornack, S. SecretSanta: flexible pipelines for functional secretome prediction. Bioinformatics 34, 2295–2296 (2018).

43. Burdukiewicz, M., Sobczyk, P., Chilimoniuk, J., Gagat, P. & Mackiewicz, P. Prediction of Signal Peptides in Proteins from Malaria Parasites. Int. J. Mol. Sci. 19, 3709 (2018).

44. Käll, L., Krogh, A. & Sonnhammer, E. L. L. A combined transmembrane topology and signal peptide prediction method. J. Mol. Biol. 338, 1027–36 (2004).

